# Out of rhythm: Compromised precision of theta-gamma coupling impairs associative memory in old age

**DOI:** 10.1101/2021.07.02.450954

**Authors:** Anna E. Karlsson, Ulman Lindenberger, Myriam C. Sander

**Affiliations:** Max Planck Institute for Human Development, Center for Lifespan Psychology, Berlin, Germany; Max Planck UCK Centre for Computational Psychiatry and Ageing Research, Berlin, Germany, and London, UK

**Keywords:** associative memory, aging, theta-gamma coupling, EEG

## Abstract

Episodic memory declines with advancing adult age. This decline is particularly pronounced when associations between items and their contexts need to be formed. According to theories of neural communication, the precise coupling of gamma power to the phase of the theta rhythm supports associative memory formation. To investigate whether age differences in associative memory are related to compromised theta–gamma coupling, we took electroencephalographic (EEG) recordings during the encoding phase of an item-context association task. Fifty-eight younger and 55 older adults studied pictures of objects superimposed on background scenes. In a recognition test, objects were presented on old or new backgrounds, and participants responded if they had seen (1) the object and (2) the object–scene pair. Theta–gamma coupling supported pair memory formation in both age groups. Whereas pair memory was associated with coupling closer to the peak of the theta rhythm, item-only memory was associated with a deviation in phase angle relative to pair memory. Furthermore, a stable relation between coupling phase and pair memory performance demonstrated that coupling closer to the peak is beneficial for associative memory. Critically, older adults’ lower pair memory was accompanied by a shift in coupling phase relative to younger adults. In concert, the present results are consistent with the hypothesis that decrements in the precision with which gamma power is coupled to the theta phase underlie the decline of associative memory in normal cognitive aging.

**Significance Statement:** According to prominent theories of neural communication, the precise coordination of oscillatory activity enables the formation of associative memories. We propose that normal cognitive aging impairs associative memory formation by compromising the precision of neural communication. We show that the coupling of high-frequency gamma power to low-frequency theta phase supports associative memory formation in both younger and older adults, with coupling closer to the theta peak benefitting memory performance. However, compared to younger adults, the coupling phase angle is shifted and more variable in older adults. We conclude that alterations in the precision of theta–gamma coupling contribute to adult age differences in associative memory.

## Introduction

The principle of Hebbian learning (1) states that memories are formed via synaptic connectivity, which is modulated by the precise timing of activity between groups of neurons (2–3). Neural oscillations reflect such rhythmic fluctuations in synchronized neural excitability, orchestrating neural communication across different temporal and spatial scales (4–7). High-frequency activity in the gamma band (~40–150 Hz) serves local processing in relatively small cortical neural assemblies over short time windows, and has been associated with the representation of item-specific information in the service of memory (8–10). In contrast, low-frequency oscillations in the theta band (~3–8 Hz) integrate information over larger temporal and spatial scales (11–12) by coordinating time windows of synaptic plasticity (13–15), and have been associated with the successful formation (16–18) and retrieval (19–21) of associative memories. Importantly, the coordination of these brain rhythms, namely the coupling of gamma power along the phase of the theta oscillation (i.e., theta–gamma cross-frequency coupling, CFC) has been demonstrated to underlie the successful formation of memories contingent upon contextual information (22–25). Thus, theta phase coding presumably provides a key mechanism for the formation of item–context associations during memory encoding (20, 26).

The decline in memory for associative information is a hallmark of normal human aging (e.g., 27–28; for reviews, see 29–31). However, surprisingly, it has not yet been investigated whether the coordination of brain rhythms underlying associative memory formation is compromised in older adults. Hence, the goal of this study was twofold: First, to investigate whether theta–gamma coupling specifically supports the successful formation of item–context associations (as compared to item-only recognition), and second, whether age-related declines in associative memory are associated with less precise theta–gamma synchronization during memory formation.

To this end, electroencephalograms (EEG) were recorded while younger and older adults engaged in an item–context association task (Fig.1) that involved the formation of associations between objects and scenes. In a subsequent recognition memory test, old and new objects were presented on top of old or new scenes, and participants were asked to judge whether (a) the object and (b) the object–scene pair were old or new. Importantly, old objects were presented either superimposed on the identical scene from encoding (match condition) or on a familiar, old scene that had been paired with other objects during encoding (mismatch condition). This design ensures that correct responses in the associative condition rely on the retrieval of the *association* between objects and scenes, and not simply on a feeling of familiarity for the respective constituents of the association. Therefore, our design allowed us to clearly separate mechanisms of item recognition from those underlying associative memory.

**Figure 1.**
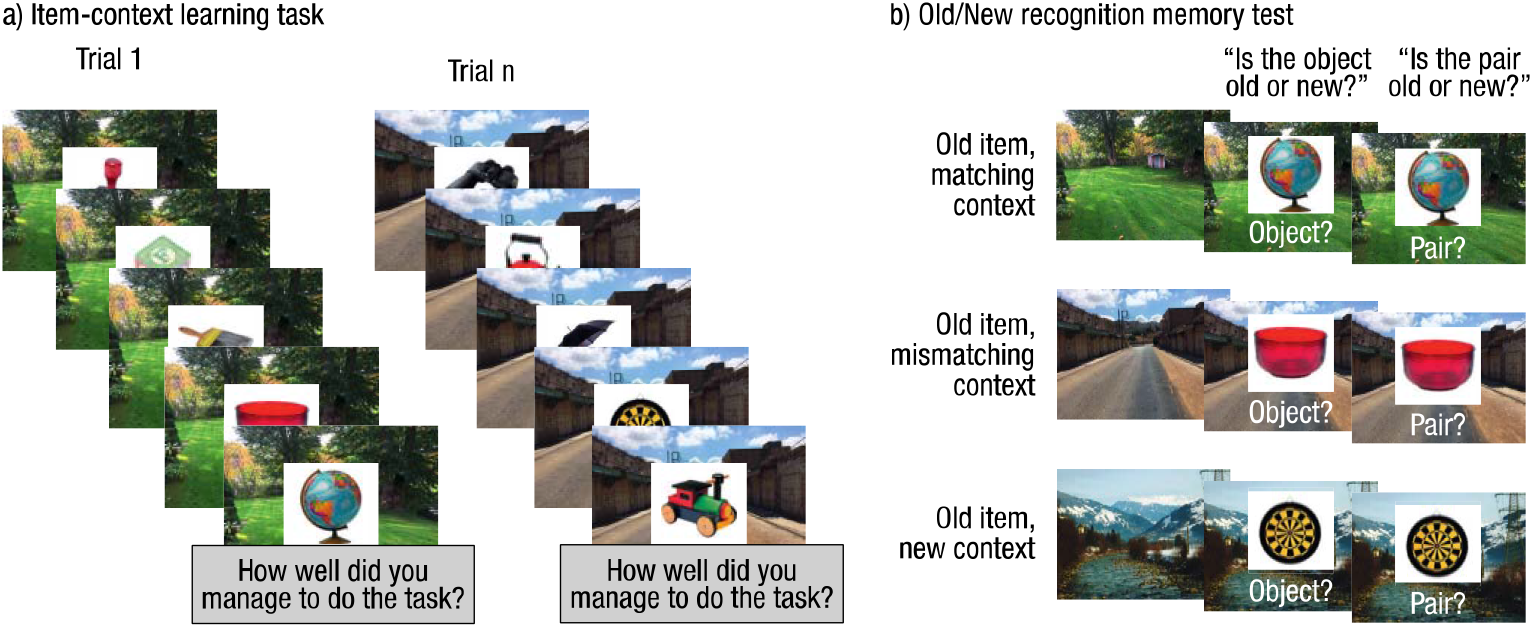
Experimental paradigm. a) During learning, participants were presented with objects superimposed on pictures of outdoor scenes and instructed to imagine using the object in the place depicted. b) Subsequently, in a surprise recognition memory test, old and new objects were presented on old (matching/mismatching) or new scenes. Participants judged first whether they had seen the object before and then whether they had seen the specific object–scene pair before. Matching scenes corresponded to the scene on which the object was presented during encoding. Mismatching scenes corresponded to old scenes from encoding that were previously paired with another object. The mismatch condition emphasized the need to base the pair memory judgement on the association between the item and the context, and not simply on a feeling of familiarity for the constituent elements of the pair.

As postulated by theory, we found that successful associative memory formation, that is, the successful subsequent retrieval of the item–context association, was preceded by a coupling of gamma power to the peak of the theta rhythm during encoding. Concurrently, item recognition was characterized by a shift in coupling phase, away from the peak, in both younger and older adults. Importantly, participants who showed a coupling closer to the theta peak also showed better associative memory. Finally, older adults’ lower pair memory performance was accompanied by a temporal shift in coupling phase compared to younger adults. Together, the present results are consistent with the hypothesis that decrements in the precision with which gamma power is coupled to the theta phase underlie the decline of associative memory in normal cognitive aging.

## Results

### Older adults show reduced pair memory performance

Item and pair memory performance were quantified by corrected recognition scores. Corrected item recognition scores were obtained by subtracting the proportion of “old item” responses to new objects from the proportion of “old item” responses to old objects on matching and mismatching scenes. Corrected pair recognition scores were obtained by subtracting the proportion of “old pair” responses to new object pairs from the proportion of correct pair responses (i.e., responding “old pair” to an old object on a matching scene and “new pair” to an old object on a mismatching scene). To investigate whether age differences are greater for pair as compared to item memory performance, corrected recognition scores were investigated in a mixed ANOVA with age (younger/older) as between- and condition (item/pair) as within-subject factors. The analyses demonstrated a main effect of condition (F(1, 111) = 412.72, *p* < .001), reflecting generally better item (*M* = 0.63, *SD* = 0.19) than pair memory performance (*M* = 0.41, *SD* = 0.14) in both age groups, and a main effect of age (F(1, 111) = 5.55, *p* = .020). In addition, a significant age-by-condition interaction (F(1, 111) = 5.06, *p* = .026) followed by pairwise comparisons showed that item memory performance did not differ between age groups (*p* > .3) while pair memory performance was significantly (*t*(94.53) = 3.53, *p* < .001) lower in older (*M*=0.36, *SD*=0.09) than in younger (*M* = 0.45, *SD* = 0.16; Fig. 2a) adults. Finally, in order to isolate explicit associative memory, we investigated pair false alarms (i.e. responding “old pair”, [FAs]) to mismatching pairs, given that this condition depends on the explicit retrieval of object-scene associations. Indeed, older adults (*M* = 0.41, *SD* = 0.21) showed a higher false alarm rate than younger adults did (*M* = 0.32, *SD* = 0.13; *t*(89.0) = −2.98, *p* = .004; Fig. 2b). Thus, we replicated the finding that item recognition remains fairly intact in old age, whereas associative memory is compromised relative to younger adults.

**Figure 2.**
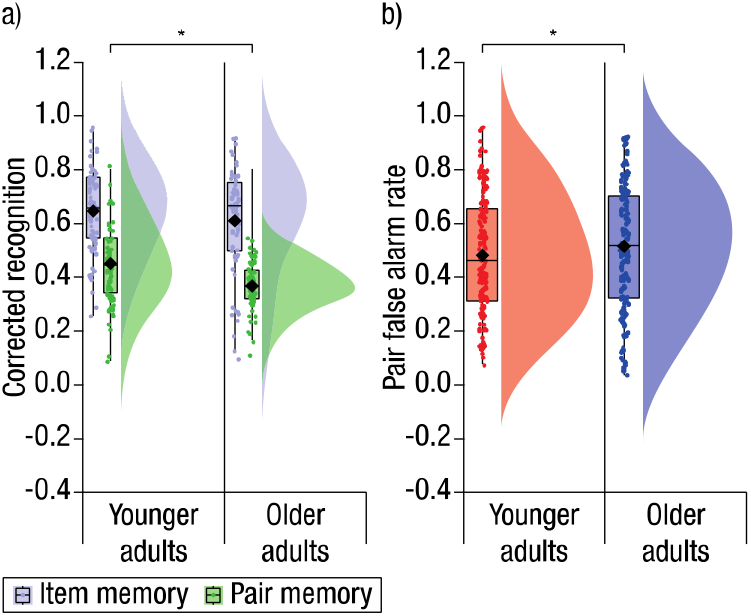
a) Corrected recognition scores (y-axis) for item (purple) and pair memory (green), demonstrating comparable item memory performance across age groups (x-axis) but reduced pair memory in older compared to younger adults. b) Pair FA rates (y-axis) demonstrate that older adults (blue) are more likely to incorrectly endorse old items on mismatching scenes as old pairs in comparison to younger adults (red). Box plots represent the interquartile range (first and third quantile), with dots representing individual participants. The rhombus indicates the mean, the horizontal bar indicates the median, and half-violin plots illustrate the sample density.

### Clear evidence for oscillatory theta activity in younger and older adults

A prerequisite for any meaningful interpretation of phase information is the actual presence of an underlying oscillation (32). Therefore, in a first step, we established the presence of oscillatory activity in the EEG signal on the single-trial level using an extension of the better oscillation detection (BOSC) method (33). This method is particularly well suited for identifying rhythmic segments in the theta frequency range on the single-trial level (see Methods). For single-trial time windows that contained theta oscillations according to BOSC, all individuals in both age groups (see Fig. S1) showed reliable peaks in the power spectrum within the theta frequency range, providing clear evidence that oscillatory theta activity was present in both age groups (for group averages of the power spectrum, see Fig. 3a). Given that the frequency of theta varies across individuals (34), we estimated the individual theta peak frequency for each participant (on rhythmic data as defined by BOSC) and used it as the phase-carrying frequency in the following analyses (see Methods). For subsequent analyses, the data were sorted into rhythmic and non-rhythmic segments (1 s) within channels and trials based on the time windows identified as oscillatory by the BOSC algorithm (see Methods for details). Note that we only included segments within a posterior cluster of channels defined as the gamma region-and-frequency of interest (see below and Fig. 3c). In order to assess differences in oscillatory activity related to item and pair memory formation, we sorted the rhythmic data into two conditions (similar to a subsequent memory analysis, e.g., 35): Rhythmic segments belonging to trials for which only the object was later recognized without memory for the object–scene association (i.e., correct item responses followed by an incorrect pair response to objects on matching or mismatching scenes; henceforth item-only memory) provided data for the item recognition condition, and rhythmic segments belonging to trials for which the object–scene association was remembered (i.e., correct item responses followed by a correct pair response to old objects on matching or mismatching scenes; henceforth pair memory) provided data for the associative memory condition. Prior to investigating the interaction between activity in the theta and gamma bands, we tested whether the temporal duration of rhythmic theta activity present during encoding differentiated between subsequent item-only memory and pair memory, and contributed to age differences therein. Based on the BOSC method, we quantified temporal duration as abundance (see 33), that is, the duration of a rhythm relative to the full time window analyzed (3 s). A mixed ANOVA with age group (younger/older) as between- and condition (item/pair) as within-subject factors showed no significant main effects or interactions (*p*s > .3; Fig. 3b). Specifically, we did not observe age-related reductions in rhythmic theta activity per se, which means that results regarding item recognition versus associative memory were not influenced by the mere occurrence of rhythmic activity.

**Figure 3.**
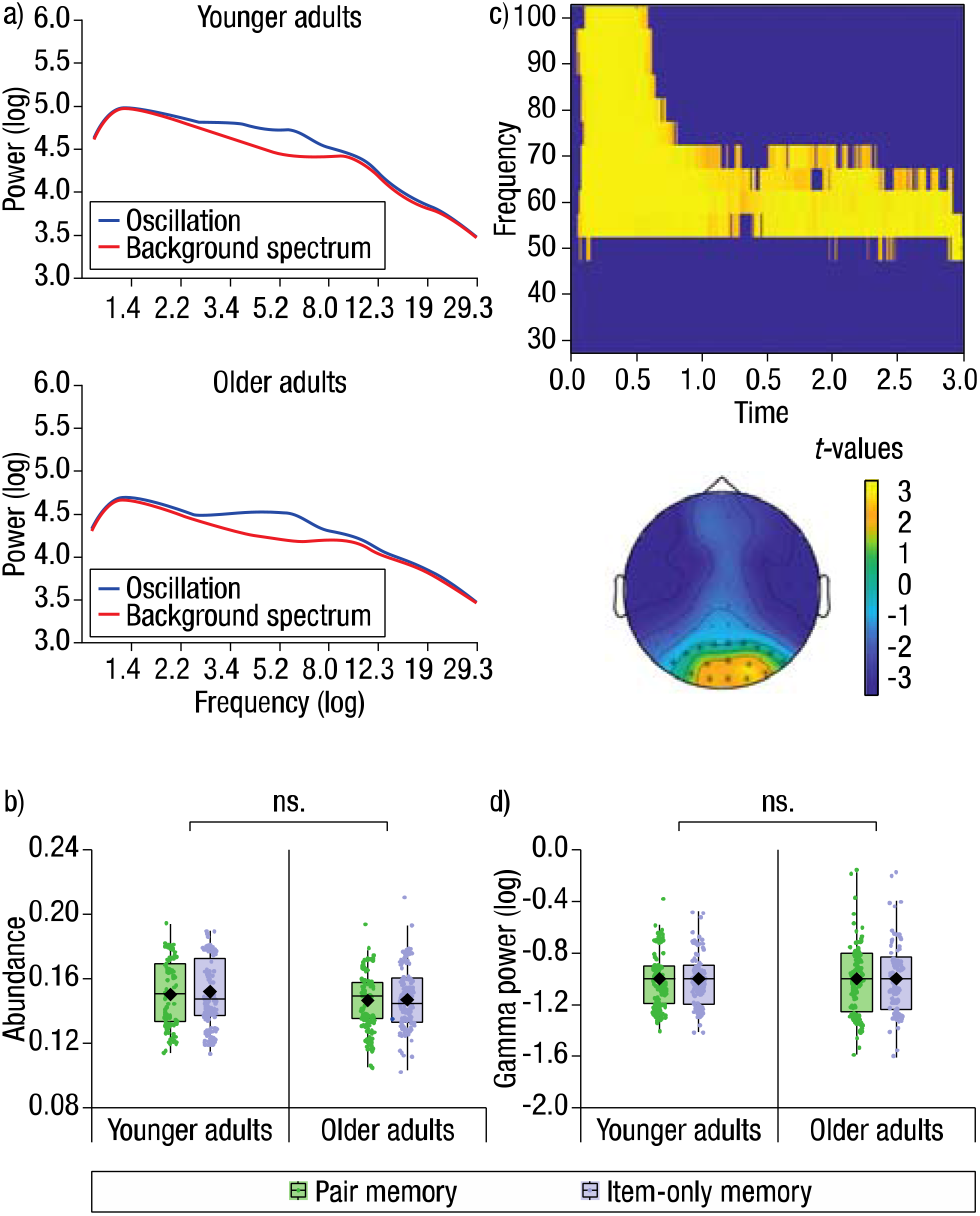
a) Younger and older adults show reliable peaks over the theta frequency range during rhythmic segments as estimated on the single-trial level. The figures illustrate power (y-axis) per frequency (x-axis), averaged over rhythmic episodes (blue) relative to the background spectrum (red). b) Illustrates the relative amount of rhythmicity over time (i.e., abundance; y-axis), which did not differ between age groups (x-axis) or between item-only memory (green) and pair memory (purple). c) *Upper*: Illustrates the identified time (x-axis) and frequency (y-axis) cluster during encoding showing reliable increases in gamma power relative to a pre-stimulus period (*t*-values). *Lower*: shows the topographical distribution of the gamma band effect averaged over time (0.4–3 s; *t*-values). Asterisks highlight significant channels. d) Gamma power (averaged across the identified cluster over rhythmic time windows) did not differ in magnitude between age groups (x-axis) or between item-only memory (green) and pair memory (purple). Box plots represent the interquartile range (first and third quantile) with dots representing individual participants. The rhombus indicates the mean and the horizontal bar indicates the median.

### Gamma power increases during memory encoding

To isolate the frequency range and topographical distribution of gamma band activity presumably representing the object information during encoding, we determined increases in neural activity during the peri-stimulus (0–3 s) period relative to the prestimulus (−0.7– −0.2 s) period using a non-parametric cluster-based permutation approach. The analysis yielded a positive effect (*p* = .009), reflecting a sustained power increase following stimulus onset (see Fig. 3c). The effect-carrying frequency range (50–75 Hz) and channel cluster (Pz, P1, P2, P3, P4, P5, P6, P7, P8, POz, PO3, PO4, PO7, PO8, O1, O2, Oz) was used as a region-and-frequency of interest for the following analyses. In analogy to the analysis on differences in theta abundance during encoding reported above, we first examined whether differences in gamma power explained subsequent memory (item vs. pair) or age group differences. Therefore, we quantified gamma power within the region-and-frequency of interest for time windows classified as rhythmic. A mixed-model ANOVA, with age (younger/older) as between- and condition (item/pair) as within-subject factors, showed no significant main effects or interactions (*p*s > .6; Fig. 3d). Thus, gamma power increases also did not show reliable differences between age groups or between subsequent item recognition and associative memory.

### Theta–gamma coupling supports associative memory formation

So far, we have demonstrated reliable oscillatory activity within the theta frequency range as well as reliable gamma power increases during encoding, but none of these factors showed differences between item and pair memory or between age groups. Next, we therefore assessed whether their interplay, namely theta–gamma CFC, would reveal differences that relate to memory performance within and between participants and age groups. To that end, we computed the mean phase angle for each participant and condition (item, pair) by weighting each theta phase bin (36 bins in total; see methods for details) with gamma power averaged over trials. Thus, an individual’s phase angle reflects the phase bin(s) at which gamma power is strongest. To test for a deviation from uniformity, which would indicate a modulation of gamma power by the theta phase, we applied the Rayleigh test on the samples of phase angles per age group and condition separately. Note that a similar analysis was performed on non-rhythmic segments, i.e. 1 second time-windows when no rhythmic theta activity was identified by BOSC, which was balanced regarding the number of trials and served as a control. The analysis demonstrated that both item-only memory and pair memory were associated with a reliable deviation from uniformity in younger (item: z = 4.12, *p* = .016; pair: z = 3.64, *p* = .026) as well as older adults (item: z = 3.65, *p* = .025; pair: z = 3.71, *p* = .024), reflecting reliable phase–amplitude coupling between theta and gamma. Importantly, non-rhythmic segments did not show a reliable deviation from uniformity in any age group (younger: z = 0.30, *p* = .74; older: z = 0.78, *p* = .46). These results suggest that gamma power increases tend to be coupled to a specific phase of the theta rhythm during memory encoding. As illustrated in Figure 4a, coupling appears to occur closer to the peak of the theta cycle for pair memory (younger adults: mean = 10.26° circular *SD* = 1.22; older adults: mean = 325.90°, circular *SD* = 1.22) as compared to item-only memory (younger adults: mean = 26.11°, circular *SD* = 1.21; older adults: mean =294.52°, circular *SD* = 1.22) in both age groups (see Table S1 for additional descriptive circular statistics). To investigate whether item-only memory, as characterized by the failure to retrieve the associated context, was associated with a deviation from the coupling phase that seemed to be optimal for successful pair memory formation, we computed the within-person absolute difference in coupling phase between item-only memory and pair memory for younger (mean = 90.53°, circular *SD* = 0.61) and older (mean = 80.55°, circular *SD* = 0.64) adults. To assess the reliability of the deviation, a phase deviation score was computed by dividing the absolute phase difference with 180 (i.e. the maximum possible difference). The Wilcoxon signed rank test demonstrated a reliable deviation of the coupling phase in both younger adults (W = 1711, z = 6.92, α = 0.05, *p* < .001) and older adults (W = 1540, z = 6.45, α = 0.05, *p* < .001), and the Wilcoxon rank sum test showed that the deviation was comparable across age groups (W = 3473, z = 0.96, α = 0.05, *p* > .3; see Fig. 4c). Thus, in agreement with our expectations, these results demonstrate that the modulation of gamma power by the phase of the theta rhythm is specific to oscillatory time-windows. Importantly, reliable theta–gamma coupling was evident in both age groups, indicating that this key mechanism of associative memory formation continues to be in effect in old age. Furthermore, in line with previous work (36), we show that there is an optimal theta phase for the formation of item–context associations, which is reflected in gamma power increases close to the peak of the theta oscillation in both younger and older adults. In contrast, item-only memory was characterized by more off-peak power increases in both age groups.

**Figure 4.**
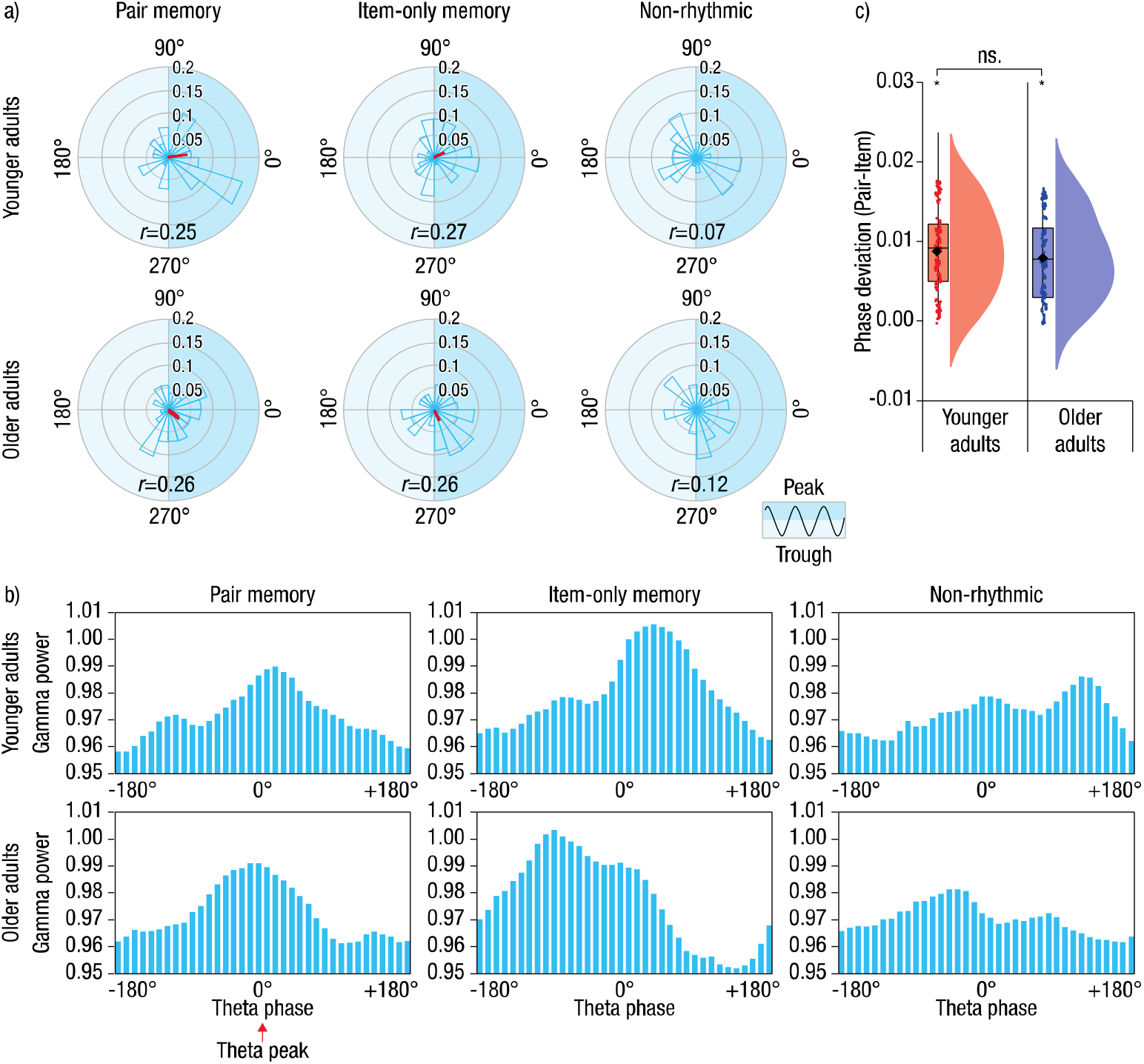
a) Angular histograms of the theta–gamma coupling phase across conditions for younger (upper) and older adults (lower). The peak of the theta cycle corresponds to phase 0° (cosine) and red lines indicate the mean direction (i.e., phase angle) and magnitude (i.e., length) of the mean resultant vector. The axis rings illustrate the proportion of samples in the histogram falling in each phase bin. Note that the red line (i.e., the vector length) has been scaled accordingly, and *r* indicates the true vector length. b) Illustrates the corresponding distributions of gamma power (y-axis) over theta phase bins (x-axis). Similarly, the peak (highlighted with red arrow) corresponds to phase 0°. c) Shows the phase deviation for younger (red) and older (blue) adults. Box plots represent the interquartile range (first and third quantile) with dots representing individual participants. The rhombus indicates the mean and the horizontal bar indicates the median. Together, the plots illustrate the presence of theta–gamma coupling during item-only memory and pair memory formation, but not during non-rhythmic segments (a– b), and the age-invariant deviation in coupling phase for item-only memory as compared to pair memory (c).

### Coupling during associative memory formation is shifted in time on older adults

To answer the key question whether aging is associated with alterations in the precision of theta–gamma coupling, that is, whether observed gamma power increases are coupled to a different phase of the theta cycle in older than in younger adults, we contrasted the phase angles between age groups using the Watson-Williams test. The analysis demonstrated significant differences in phase angles between age groups for pair (F(1,112) = 4.81, *p* = .030) and for item-only (F(1,112) = 20.13, *p* < .001) memory. Thus, although both age groups tended to couple close to the peak of the theta oscillation, there was a clear temporal shift in coupling phase in older adults relative to younger adults. Interestingly, this age-related shift in coupling phase was evident for both subsequent item-only memory and pair memory (see Fig. 4a for an age comparison of the coupling phase directions and Fig. 5a for an age comparison of the gamma power distributions). As illustrated in Figure 5b, younger adults showed gamma power increases more consistently around the peak of the theta rhythm while older adults showed a more distributed coupling pattern. Finally, to assess whether inter-individual differences in coupling phase (that is, the theta phase at which gamma power increases were most prominent) were related to inter-individual differences in pair memory performance, we computed the linear-circular correlation between the individual phase angle within pair memory trials and the proportion of correct pair responses. The analysis demonstrated a positive relationship, indicating that coupling closer to the peak of the theta oscillation was beneficial for associative memory (*r* = 0.31, *p* = .005; Fig. 5c). Note that the correlation remained fairly reliable within each age group (younger: r = 0.31, *p* = .057; older: r = 0.35, *p* = .037), suggesting that the relationship is not driven solely by one age group. To further understand this relationship, a median split was performed on the proportion of correct pair responses per each age group, and theta–gamma coupling during pair memory formation was assessed for high- and low-performing younger and older adults separately. The Rayleigh test demonstrated that high-performing younger adults (z = 6.52, *p* = .001; mean = 346.26°, circular *SD* = 1.03) and older adults (z = 4.11, *p* = .02; mean = 346.62°, circular *SD* = 1.12) showed reliable theta–gamma coupling, while neither age group of low-performers did (*p*s > .29; see Table S1 for additional descriptive circular statistics). Further, the Watson-Williams test (F(1,57) = 0.00, *p* = .99) demonstrated that gamma power in high-performing older adults’ were as precisely coupled to the peak of the theta rhythm as in high-performing younger adults. These findings provide strong evidence for the claim that the coupling of gamma power to theta phase during encoding supports associative memory formation. In addition, the results are consistent with the hypothesis that adult age differences in associative memory reflect a decline in the precision of this coupling mechanism.

**Figure 5.**
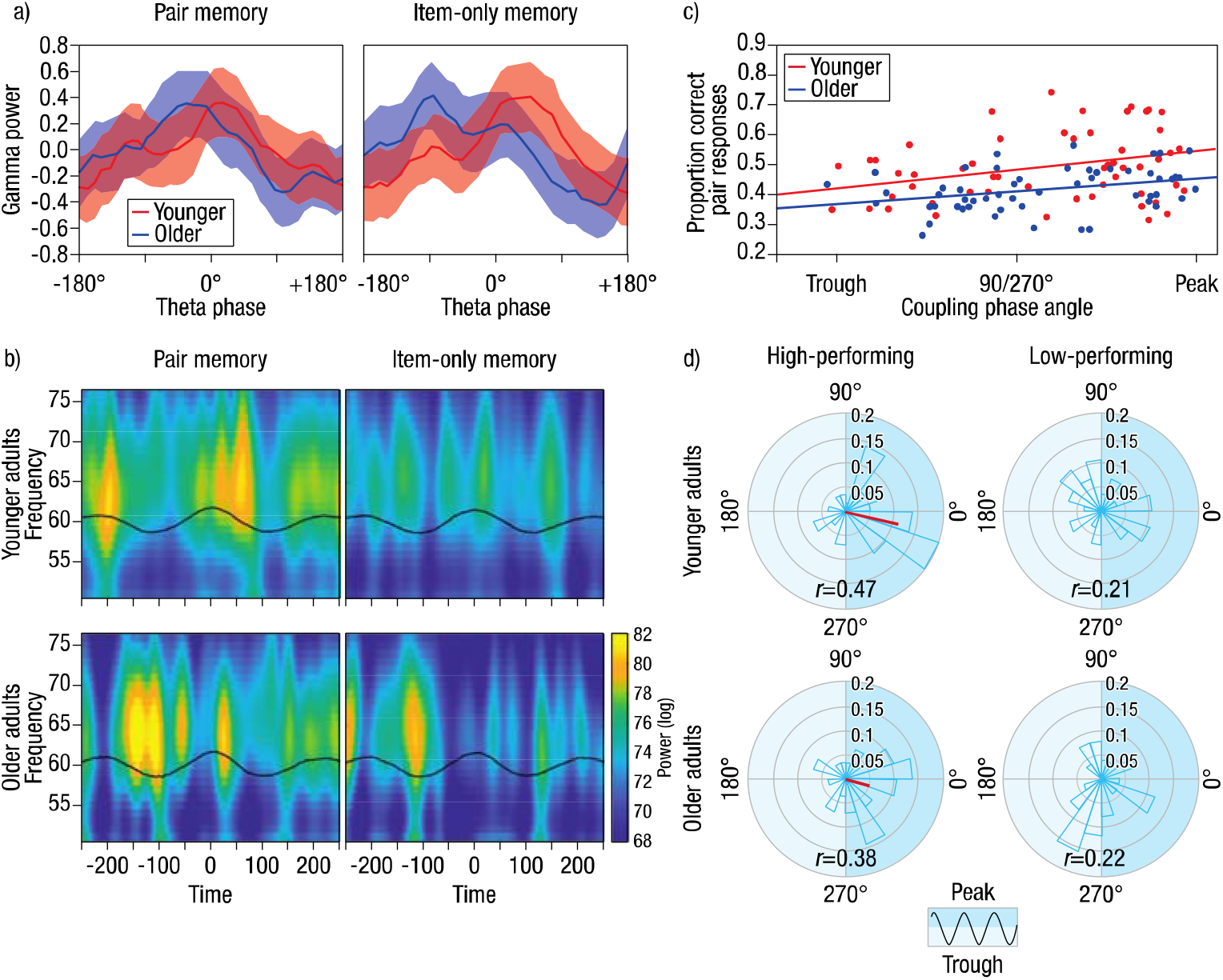
a) Illustration of age differences in the distribution of gamma power (*y*-axis) over the theta phase bins (x-axis) for pair memory (left) and item-only memory (right). Shaded area indicates 95% confidence interval and phase 0° corresponds to the theta peak (cosine). The plots illustrate the shift in coupling phase for older as compared to younger adults. b) Illustration of the time (x-axis) and frequency (*y*-axis) power spectrum within the gamma band (50–75 Hz) for pair memory (left) and item-only memory (right). The black line represents the unfiltered event-related potential (ERP). The data were aligned to the middle peak (time = 0) of the theta rhythm. The plots illustrate the decreased coupling precision in older as compared to younger adults. c) The relationship between the proportion of correct pair responses (y-axis) and individual coupling phase (x-axis; angle in degrees). For illustrative purposes only, absolute phase angles are plotted. Colored lines and dots indicate regression lines for the two age groups and individual data points respectively. The plot demonstrates a positive association, suggesting that coupling around the peak is beneficial for pair memory performance. d) Angular histograms demonstrating the greater coupling precision evident in high-performing, as compared to low-performing, younger and older adults. The peak of the theta cycle corresponds to phase 0° (cosine) and red lines indicate the mean direction (i.e., phase angle) and magnitude (i.e., length) of the mean resultant vector. The axis rings illustrate the proportion of samples in the histogram falling in each phase bin. Note that the red line (i.e., the vector length) has been scaled accordingly, and *r* indicates the true vector length.

## Discussion

Accumulating evidence suggests that the precise coupling between gamma power increases and the phase of rhythmic theta activation (e.g., 10, 25) supports the formation of associations between different elements of an event, an essential feature of episodic memory (37). Normal human aging is associated with declines in episodic memory, and age differences are greatest for memory for associative information (e.g., 29). Accordingly, it has been proposed that aging is associated with reductions in the ability to associate distinct elements of an episode into a coherent memory representation during encoding (38). However, the question whether normal aging compromises the precision of theta–gamma coupling during the formation of associative memories has not been addressed thus far.

Here, we showed that theta–gamma CFC supports successful memory formation in an item–context association task in both younger and older adults. First, we documented the presence of oscillatory activity in the theta frequency range in both younger and older adults (Fig. 3a–b). Similarly, the increase in gamma power during encoding did not differ by age group (Fig. 3c–d). We then provided evidence for the key role of theta– gamma CFC in associative memory formation by showing that gamma power is coupled to the phase of the theta rhythm during memory encoding. Theta–gamma coupling was evident irrespective of whether only the item or the pair were correctly recognized at test. However, the coupling phase of trials in which later only the object was recognized (without its associated scene, or item-only memory) deviated from the coupling phase of trials in which the object–scene pair (that is, the association of the object with its scene, or pair memory) was successfully retrieved (Fig. 4). In comparison to item-only memory formation, pair memory formation was characterized by gamma power increases that were closer to the peak of theta oscillation – presumably the “optimal” phase for memory formation (13, 36). Importantly, this effect was present in both younger and older adults. Furthermore, individual differences in associative memory were correlated with individual differences in the theta coupling phase (Fig. 5c) in both age groups. Critically, we observed reliable differences in coupling phase between age groups, with older adults’ lower pair memory performance being associated with a phase shift away from the theta peak (Fig. 5a–b; see also Fig. 4a). Taken together, our results support the hypothesis that normal aging compromises the precision of neural communication during the formation of associative memories.

### Theta–gamma coupling supports the formation of item–context associations

The nesting of gamma band activity within the theta cycle reflects the synchronization of distributed neural populations, whereby the phase of the theta cycle determines time windows of synaptic plasticity, inducing long-term potentiation and long-term depression (LTP and LTD; e.g., 14–15). Accordingly, leading theories on the role of a theta–gamma code in memory (10, 13, 39) posit that the theta rhythm regulates the precise timing of neural activity between different cell populations, and thereby promotes the strengthening of synaptic connections that serves associative memory formation. Recent work has demonstrated that theta–gamma CFC supports episodic memory contingent upon contextual information (22–25, 40). However, most studies did not explicitly assess associative memory but used confidence ratings (24) or the remember/know procedure (40–41) to infer recollection at the time of testing. In the present design, pair memory was explicitly tested, allowing us to disentangle potential differences in CFC underlying subsequent item-only memory versus pair memory. We found that item-only versus pair memory were associated with different coupling patterns. In particular, coupling closer to the peak of the theta rhythm was beneficial for later pair memory. In contrast, when items were later recognized without their associated context, we observed a reliable deviation of ~80–90° from the coupling phase optimal for successful associative memory (Fig. 4). This finding suggests that the failure to subsequently retrieve the association between an item and its context is associated with gamma power increases occurring at a substantially different theta phase that is shifted away from the theta peak. Thus, our results extend previous findings by demonstrating that theta–gamma CFC during encoding specifically supports the formation of item–context *associations*. The finding that neither the temporal duration of rhythmicity in the theta band (Fig. 3b) nor the strength of cortical activation in the gamma band (Fig. 3d) present at encoding distinguished between subsequent item-only memory and pair memory substantiates the proposal that associative memory formation is dependent upon the precise timing of the interaction between theta and gamma activity. Thus, we argue that cortical representations of item and context information, as reflected in activity in the gamma band, are bound via the theta rhythm (16, 20), which assure the exact timing in neural communication to lay down new associative memories.

Most prior observations of theta–gamma CFC during memory formation have been made in the hippocampus and/or the surrounding medial temporal lobe (MTL), an associative hub strongly implicated in episodic memory formation (42–44) where theta is the dominant rhythm (45). The extent to which scalp-recorded theta reflects hippocampally versus neocortically generated theta is not fully understood. It has been suggested that cortical theta is driven by hippocampal projections (46), and that synchronized gamma activity in neural assemblies across distributed cortical regions are biased by the hippocampal theta phase (12). In addition, increased phase coherence in the theta band has been demonstrated between frontal and posterior cortical networks during the formation of item–context associations (47–48, see 11 for review), and between sub-regions within the MTL in support of successful memory formation (49). Moreover, Clouter and colleagues (16) showed that inducing auditory and visual oscillatory entrainment to a synchronized theta phase (i.e., with zero phase lag) was beneficial for subsequent associative memory as compared to a desynchronized entrainment phase (i.e., a phase lag of 90° or more). Taken together, these results provide strong evidence that the theta rhythm binds together distinct pieces of information via phase synchronization, both locally and across larger spatial distances, allowing associative information to be coded by the hippocampal network. Given that the task used in the present study can be assumed to rely heavily on hippocampally mediated associative processes, it is likely that the theta–gamma coupling observed in the present study is triggered by or at least interacting with the hippocampal network in coordinating the formation of associations between item and context information.

### Lower pair memory in older adults is accompanied by a temporal shift in theta– gamma coupling

The associative deficit hypothesis posits that age-related declines in episodic memory reflect difficulties in forming associations between distinct elements of an episode (38). Accordingly, we found that item recognition was comparable across age groups, whereas older adults showed reduced memory for associative information relative to younger adults (Fig. 2). These age differences in pair memory performance were accompanied by age differences in coupling phase. While coupling around the peak of the theta rhythm was beneficial for pair memory formation in both age groups, older adults showed a reliable shift in coupling phase relative to younger adults for both subsequent pair memory and item-only memory (i.e., objects later recognized with and without the associated scene; Figs. 4a–b, 5a–b). Hence, as a group, older adults consistently showed less precise coupling to the optimal phase of the theta rhythm than young adults. Importantly, the observation that neither the temporal duration of rhythmicity nor gamma power increases during encoding differed between age groups further corroborates the proposal that the reduction seen in associative memory in the older age group was due to an alteration of the precise interaction between theta and gamma activity.

It may surprise that age differences in coupling phase were present regardless of subsequent associative memory success. However, trials here defined as item-only memory included correct item responses *followed by incorrect pair responses*, thus, trials in which associative information was not available or not correctly retrieved. The proportion of these trials differed between age groups, as reflected in higher false alarm rates to mismatching pairs in older as compared to younger adults. Higher occurrence of these false alarms, that is, false recognition of re-arranged pairs, is a common finding in aging research and has been related to age differences in hippocampal activation (50–51) and to the volume of a subregion of the hippocampus (DG/CA3; 52) that supports episodic memory formation (53). Accordingly, the observed age differences in CFC for item-only memory likely reflect the increased rates of false alarms in older adults resulting from an attenuated coding of the association by the hippocampal system.

Our results add to previous findings indicating that disruptions in the dynamics of neural communication underlie senescent cognitive changes (54). Previously, decreased synchrony in the theta band between frontal and posterior regions (55), and altered synchronization between theta and gamma activity (56; see also 57) has been associated with reduced working memory performance in older adults. The present results extend these observations of compromised neural communication as a marker of age-related cognitive decline to the domain of episodic memory formation. At the same time, it is important to note that those older adults who showed a more “youth-like” theta–gamma coupling pattern also showed better associative memory, as revealed by a stable relationship between individual coupling phase and the proportion of correct pair responses (cf. 58–60). In fact, among high-performing older adults, coupling occurred as precisely to the peak of the theta rhythm as among high-performing younger adults. This is in line with the proposal that cognitive performance in old age is determined by the level of preservation of neural structures and functionally specialized processes (60–61, 30). In particular, maintenance of hippocampal structure may preserve hippocampally mediated functions such as the precise timing of neural synchronization in high performing older adults. Thus, while the temporal dispersion in theta–gamma CFC contributes to age-related declines in associative episodic memory, older adults who are able to preserve a youth-like coupling precision also maintain their episodic memory function to a greater extent. However, when the neural system fails to consistently couple to the optimal phase of the theta rhythm, the induction of synaptic plasticity is attenuated, and the coding of associative information by the MTL is compromised, resulting in reductions of associative memory performance with advancing adult age.

Finally, it is noteworthy that neurotransmitters, e.g. dopamine and norepinephrine, have been shown to modulate neural plasticity (for reviews, see 62–63), and recent work suggests that dopamine may modulate the coupling between higher and lower frequencies (64). In addition, individual differences in episodic memory have been related to differences in neurotransmitter function (65, for review, see 66). Thus, individual differences in the stability of the underlying neurochemical system may be a contributing factor to the observed differences in coupling precision. Future work may focus on the link between age-related changes in neuromodulating transmitters and theta-gamma coupling, and the relation to hippocampus-dependent associative processes.

## Conclusion

Prominent neurocognitive theories of memory have for long suggested that rhythmic modulations of synchronized neural excitability serve neural communication, and that reductions in the precision of cross-frequency coupling come at a cost for cognitive performance. Here we show that during the formation of item–context associations, gamma power increases are coupled to an underlying theta rhythm in both younger and older adults. Coupling closer to the peak of the theta rhythm is associated with better pair memory, pointing to a phase angle that is optimal for facilitating associative binding. Older adults showed lower pair memory performance and a large deviation from younger adults’ mean phase direction. At the same time, high-performing older adults showed more youth-like patterns of theta–gamma coupling. The present pattern of findings is consistent with the hypothesis that normal aging reduces the precision of neural communication, leading to age-related impairments in the formation of episodic memories.

## Materials and Methods

### Participants

Younger (*n* = 65) and older (*n* = 74) healthy, right-handed, German-speaking adults were recruited via a database of the Max Planck Institute for Human Development to participate in the study. They received compensation of €12/hour. A total of 24 participants were excluded due to technical problems (6), drop out (3), noisy EEG data (1), memory performance below chance (2; see Behavioral analysis), and due to too few trials per condition (12; see EEG recording and preprocessing). The remaining sample consisted of 58 younger (33 females, *M_age_* = 25.0, *SD_age_* = 3.1, range 20–31 years) and 55 older adults (24 females, *M*_age_ = 69.6, *SD*_age_ = 3.7, range 64–76 years). To further characterize the sample, all participants completed a demographic questionnaire and additional cognitive tests. In addition, older adults were assessed using the Mini Mental State Examination (MMST; 67). A MMST score of 27 or higher indicates normal cognition, whereas a MMST score of 19–23 indicates mild cognitive impairment (68). Performance in these tasks (see Table S2) was similar to other cognitive neuroscience studies previously run at our research center (e.g., 69–71) and showed a typical pattern of age differences. The ethics committee of the Deutsche Gesellschaft für Psychologie (DGPs) approved the study.

### Stimuli

The experiment was programmed in MATLAB (version 2016b; MathWorks Inc., Natick, MA, USA), using the Psychophysics Toolbox (72). The stimulus pool consisted of 498 colored pictures (400 x 400 pixels) of everyday objects (e.g., ball, teapot) and 210 colored pictures (1280 x 960 pixels) of outdoor scenes (e.g., forest, beach).

### Experimental paradigm

The experiment consisted of four parts, a pre-learning phase, a learning phase, a postlearning phase, and a retrieval phase, completed during one day with short breaks in between the sessions, including a 40-minute lunch break after the learning phase. For the purpose of the present analysis, we focus on the learning and retrieval phases. However, note that the objects and scenes presented in the learning phase, and thus serving as old in the retrieval phase, had been shown in a target detection task during the pre-learning phase performed in the MRT. Furthermore, the objects were also presented in the post-learning phase in a similar target detection task.

The learning phase was performed in a dimly lit room that was electromagnetically and acoustically shielded. Participants’ neural activity was recorded with EEG and their eye movements were monitored with an eye-tracker. Prior to the task, participants received instructions and completed two short practice rounds that were repeated if necessary. The experimental session started with written instructions on the screen and the experimenter initiated the session with a button press.

During encoding, 250 randomly drawn objects and 50 randomly drawn scenes were presented together in an item–context association task. Each trial started with a jittered fixation cross (~1–1.5 s), followed by the presentation of a scene. Next, five objects were presented (3 s each) sequentially superimposed on the center of the scene, and each presentation was separated by a fixation cross (2 s). To ensure explicit attempts to associate objects and scenes and promote binding in both age groups (e.g., 73–74), participants were instructed to imagine using the presented object in the place depicted in the scene. After the presentation of the five objects, participants indicated on a three-point Likert scale how well they managed to do the task.

At test, participants were presented with the old objects intermixed with 150 randomly drawn new objects, presented on either a new (*n* = 100) or an old scene (*n* = 50). This resulted in five test conditions: old objects presented on their original scene from study (match condition, *n* = 100 trials), old objects on old but mismatching scenes from study (mismatch condition, *n* = 100 trials), old objects on new scenes (old-new condition, *n* = 50 trials), new objects on old scenes from study (new-old condition, *n* =100 trials) and new objects on new scenes (new-new condition, *n* = 50 trials). We constrained the occurrence of old objects and scenes in the recognition test in several ways: (1) The five objects belonging to the same learning sequence (i.e., presented on the same scene) were assigned to the three test conditions involving old objects as follows: Two objects were assigned to the match condition, two objects to the mismatch condition, and one object to the old-new condition. Since the position of the object within the learning sequence may influence memory performance, we made sure that each condition contained an equal number of objects from each encoding position. (2) The order of trials was randomized, with the constraint that no more than three consecutive trials could stem from the same condition. (3) Each scene from encoding was presented a total of five times (within the different test conditions, see above), but never in consecutive trials during tests.

At test, each trial started with a fixation cross (0.5 s) followed by the presentation of a scene (1 s). Next, an object was presented superimposed on the center of the scene. Participants were first required to respond whether the object was old or new (max 3 s) and subsequently whether the specific object–scene pair was old or new (max 4 s). Responses were recorded using a response box and the mapping of button to response (“old”/“new”) was counterbalanced across participants.

### Behavioral analysis

Behavioral responses were analyzed as item and pair responses. First, to assess item chance-level performance, the proportion of correct item responses was calculated across all three old item conditions (match, mismatch, old-new) with the chance level set to 0.35 (derived from the multiplication of the response probability by the proportion of trials with an old item, i.e., 0.5 x 250/400 = 0.31). Participants with performance below this chance level were excluded from the analysis. In order to control for response biases, item and pair memory performance were quantified as corrected recognition scores. Corrected item recognition scores were obtained by subtracting the proportion of “old item” responses to new objects from the proportion of “old item” responses to old objects (from the match and mismatch conditions only). Similarly, corrected pair recognition scores were obtained by subtracting the proportion of “old pair” responses to new objects from the proportion of correct pair responses (i.e., responding “old pair” to an old object on a matching scene or “new pair” to an old object on a mismatching scene; 75). To assess age differences in item and pair memory performance, corrected recognition scores were analyzed in a mixed effects ANOVA, with age (younger/older) as between- and condition (item/pair) as within-subjects factor. Any significant interactions were followed up by pairwise comparisons. Finally, since correctly rejecting an old object with a mismatching scene as a new pair depends on the explicit retrieval of the association, and cannot rely simply on memory for the constituent parts, associative memory is most clearly emphasized in this condition. Therefore, to assess explicit associative memory and age differences therein, we compared pair false alarm rates (i.e., “old pair” responses) to old objects on mismatching scenes between age groups using a *t*-test. The data was analysed using R 3.5.2 (R Development Core Team, 2018).

### EEG recording and preprocessing

The EEG was recorded with BrainVision Recorder (Brain Vision Products GmbH, Gilching, Germany) from a 61 Ag/Ag-Cl electrodes embedded cap, which was placed according to the 10-10 system (1000 Hz sampling rate; right mastoid reference). An electrode above the forehead (AFz) served as ground. To measure eye movements with the electro-oculogram (EOG), electrodes were placed below the left eye and at the left and right outer canthi. Electrode impedances were kept below 5 kΩ during the EEG measurement. In addition, participants’ heartrate was recorded via electrocardiogram (ECG) to remove possible cardiovascular artifacts from the EEG signal (see below). EEG pre-processing was performed using FieldTrip (76), EEGLAB (77), and custom-written MatLab code (MathWorks Inc., Natick, MA). Before preprocessing, eye movement and EEG data were merged along the time vectors to ensure equal time-point zero across modalities. The EEG data were filtered (4th filter order) with a bandpass of 1–150 Hz and re-referenced to the linked mastoid channels. The ECG data were filtered with a band-stop filter (48–52 Hz) and appended to the EEG data. Next, the data were segmented into epochs of 2 s, and each trial was visually inspected. Epochs containing strong artifacts not related to eye movements or blinks were temporarily excluded for a following independent component analysis (ICA). In addition, any channel strongly contaminated by artifacts was excluded. Blink, eye-movement, muscle, and heartbeat artifacts were detected using ICA (78) and removed from the signal. In addition, saccade-related transient spike potentials were identified using the COSTRAP algorithm and removed from the signal as independent components (79). Artifact-contaminated channels and trials (determined across all epochs) were automatically identified (1) using the FASTER algorithm (80) and (2) by detecting outliers exceeding four standard deviations of the kurtosis of the distribution of power values in each epoch within low- (0.2–2 Hz) or high-frequency (30–100 Hz) bands respectively. Channels labeled as artifact-contaminated were interpolated using spherical splines (81). Finally, the data were segmented into 7-s epochs, ranging from 2 s before, to 5 s after stimulus onset and sorted into conditions of interest: item-only memory (i.e., objects correctly remembered without the associated scene), and pair memory (i.e., objects and the associated scene correctly remembered). A second visual inspection of each trial per participant was performed and any remaining artifact-contaminated trials were excluded from further analysis, resulting in an average of 5.95% of trials being rejected. Participants with less than 10 trials per condition were excluded from the analyses (see Table S3 for the distribution of trials across conditions and age groups).

### Single-trial rhythm detection and individual theta peak frequency estimation

Since phase information is only meaningful in the presence of an underlying oscillation (32), we employed the extended BOSC method (33, 82–83) to identify rhythmic events in the theta frequency range (3–7 Hz) on the single-trial level. The method assumes that rhythmic events have stronger power than the 1/f background spectrum, and should be present for a predefined minimum number of cycles at a given frequency. By means of robust regression, the procedure dissociates narrowband peaks from the aperiodic background spectrum (for more details, see 33), and identifies time points per trial and channel for which the signal meets these criteria and can thus be considered to reflect an oscillation. In the present study, a three-cycle threshold was set as the temporal criterion and the 95th percentile above the individual background spectrum was defined as the power threshold. EEG data were decomposed into 49 logarithmically spaced center frequencies between 1 and 64 Hz using a 5-cycle wavelet. The most dominant peaks between 3–7 Hz (theta) and 8–15 Hz (alpha) were removed prior to fitting the 1/f slopes. Next, rhythmic segments were estimated, demonstrating reliable peaks in the theta band in all subjects in both age groups (Fig. 3a and S1). Because theta frequency varies across individuals (e.g., 34), we estimated the individual theta peak frequency to use as the phase-carrying frequency in the subsequent theta–gamma coupling analyses (for a similar approach, see 9, 23). To this end, the power spectrum was averaged across all rhythmic segments for each channel and trial. By means of local maxima, a peak frequency was estimated per channel and the final individual theta peak frequency range was defined as the average peak frequency across channels ±0.5 Hz.

Next, rhythmic segments of 1 s surrounding the middle peak of the theta rhythm, as estimated by BOSC, were extracted. Peaks were detected by means of local maxima (phase > 2Π – 0.1) and used to align the ERP and the gamma power spectrum to the time point reflecting the peak (time = 0) of the oscillation in the illustrations in Figure 5b. Note that 1-s segments were chosen to cover at least three cycles of the theta center frequency (minimum 3 Hz). In addition, the first 0.2 s following stimulus onset were excluded to avoid confounds by evoked responses. Finally, we grouped the rhythmic segments over the posterior channels defined as a region of interest (see Fig. 3c; see below) and averaged over segments within a given trial. Similarly, non-rhythmic 1-s segments were extracted from time periods not characterized as rhythmic by BOSC. To keep the number of rhythmic and non-rhythmic segments equal for later analysis, we randomly selected as many non-rhythmic as rhythmic segments from the total sample of non-rhythmic segments per channel and trial. Similarly, to balance the number of trials, we then randomly selected as many trials as the average across the two conditions (item-only and pair).

Next, we investigated whether the temporal duration of rhythmicity present during encoding differed between age groups and whether it alone already predicted subsequent item-only memory and pair memory. The temporal duration of rhythmicity present was quantified as abundance, i.e., temporal duration relative to the full time window analyzed (3 s). We conducted a mixed-effect ANOVA with age (younger/older) as between- and condition (item/pair) as within-subject factors.

### Gamma power estimation

To determine the frequency range and topographical distribution of gamma band activity involved in memory encoding, we assessed gamma power increases relative to the prestimulus time period as follows: Gamma power was estimated using multi-tapers (7; Slepian), with 0.4 s window length, a step size of 0.08 s, and a frequency resolution of 5 Hz. The power spectrum was log-transformed at the single-trial level (84). The peristimulus power spectrum (0–3 s) was then contrasted against the pre-stimulus power spectrum (−0.7 – −0.2 s), within participant. Baseline data were averaged over trials and time, and power increases were quantified by computing single sample *t*-statistics for any given channel-frequency-time point. To test statistical significance, the distribution of t-values was tested against a null distribution on the group level using a non-parametric cluster-based permutation approach (85). Initially, clusters were formed based on univariate, one-sided independent samples *t*-statistics for each channel-frequency-time point. The threshold for data points to be included in a cluster was set to *p* = .01 and the spatial constraint was set to a minimum of two neighboring channels. Next, the significance of the summed *t*-values was assessed in comparison to a permutation null distribution, obtained by switching the condition labels and re-computing the *t*-test 2000 times. The final cluster *p*-value (i.e., the Monte Carlo significance probability) is the proportion of random partitions in which the cluster-level statistics were exceeded. The threshold was set to *p*-values below .05, i.e., one-sided significance threshold. The channel-frequency cluster identified in this analysis (see Fig. 3c) was used to define a gamma-band region-and-frequency of interest for further analyses. Finally, we tested whether gamma power increases differed between age groups and between later item-only memory and pair memory. To that end, power values within the region-and-frequency of interest were sorted based on subsequent memory (item-only/pair; as described above), averaged across rhythmic segments, and analyzed in a mixed-design ANOVA with age (younger/older) as between- and condition (item/pair) as within-subject factors.

### Cross-frequency coupling

To investigate whether gamma power increases are modulated by the theta phase, we estimated the coupling between theta phase and gamma power in the following way: To derive the phase of the theta oscillation, the pre-processed signal was filtered with a two-pass bandpass filter (filter order = 3), with a bandwidth centered at the individual peak frequency range (i.e., peak frequency +/- 0.5 Hz). The analytic phase time series were estimated via Hilbert transform. To derive the instantaneous power of the modulated frequency (i.e., within in the gamma frequency range), a two-pass bandpass filter (filter order = 3) was applied from 50 to 75 Hz, comprising the frequency range showing reliable power increases during memory encoding (see Fig. 3c), in steps of 5 Hz. To avoid biased results towards slower frequencies (32), the bandwidth was set to +/- 7.5 Hz. The Hilbert-derived complex signal was z-scored within frequency bands to normalize amplitudes across frequencies, and finally squared to estimate the instantaneous power time series. Next, the power time series were separated into rhythmic and non-rhythmic segments (see section on single-trial rhythm detection) and sorted into 36 phase bins (10° bin width) according to the phase time series within a given channel and trial. To assess the presence of theta–gamma coupling, we tested for deviations from uniformity using the Rayleigh test, which evaluates whether the resultant vector length is large enough to indicate a non-uniform distribution. To this end, phase angles were computed for each participant and condition (item-only/pair) by weighting each phase bin by gamma power averaged across trials. In order to assess whether gamma power is modulated by the theta phase for subsequent item-only memory and pair memory in both age groups, the Rayleigh test was applied to each condition and age group separately. In addition, the randomly selected non-rhythmic segments were used as a control condition and analyzed in a similar fashion. The Watson-Williams test, which assesses whether the mean direction (i.e., coupling phase angle) differs between two groups, was used to assess age differences in coupling phase angles. To examine whether item-only memory is characterized by a deviation in coupling phase, the absolute difference in phase angle between item-only memory and pair memory was computed for each participant. To assess the statistical significance, a phase deviation score was computed by dividing the absolute phase difference with the maximum possible value (i.e., Pair - Item-only /180) and the Wilcoxon signed rank test was applied per each age group. To assess age differences in phase deviation, the Wilcoxon rank sum test was used to contrast the age groups. Finally, to examine the relationship between coupling phase and associative memory, a linear-circular correlation between individual phase angle and the proportion of correct pair responses was computed. To better illustrate the heterogeneity within our sample, we additionally performed a median split on the pair hit rates (within age groups) and theta–gamma coupling in low- and high-performing younger and older adults were examined by means of the Rayleigh test and age groups were contrasted using the Watson-Williams test. The analysis was performed using the CircStats toolbox (86) and custom written MATLAB code.

## Acknowledgments

Data collection and analysis was conducted within the projects Lifespan Age Differences in Memory Representations (LIME) (PI: MCS) and Lifespan Rhythms of Memory and Cognition (RHYME) (PI: MWB) at the Max Planck Institute for Human Development, Germany. AEK was a fellow of the International Max Planck Research School on the Life Course (LIFE). MCS was supported by the MINERVA program of the Max Planck Society. We thank Gabriele Faust and all student assistants who helped with organization and data collection, Julian Kosciessa for valuable discussion, Julia Delius for editorial assistance, all project members for their helpful feedback, and all study participants for their time.

## Author Contributions

A.E.K, and M.C.S. designed research; A.E.K. performed research, and analyzed the data; M.C.S., and U.L. provided funding, resources, project administration and supervision; A.E.K. wrote the paper; and A.E.K, U.L., and M.C.S. revised and edited the paper.

## Competing Interest Statement

The authors declare no competing interests.

## Data Availability Statement

Code and data will be made publicly available upon publication.

## Ethics Approval Statement

All experiments were approved by the ethics committee of the Deutsche Gesellschaft für Psychologie (DGPs).

## Supplements

**Figure S1.**
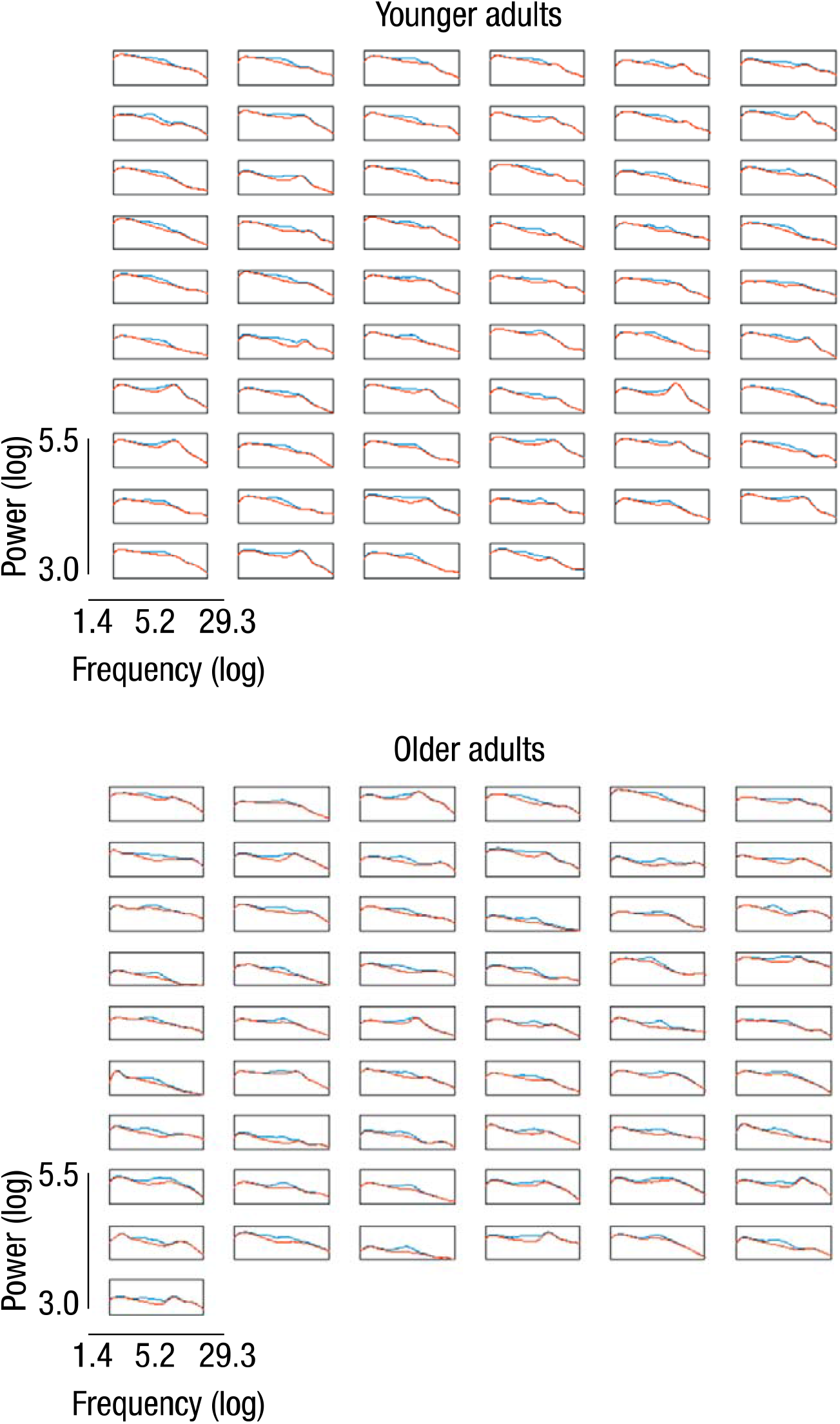
Illustrates power (y-axis) across frequencies (x-axis) for each subject by age group: younger (top) and older (bottom) adults. Power was averaged over rhythmic (blue) segments and plotted against the background spectrum (red). The plots demonstrate that each subject in both age groups showed a clear peak in the power spectrum over the theta frequency range.

**Table S1.** Descriptive circular statistics.

|  | Younger adults |  |  |  | Older adults |  |  |  |
| --- | --- | --- | --- | --- | --- | --- | --- | --- |
|  | Pair | Item | Pair – Item | High-perform. | Pair | Item | Pair – Item | High-perform. |
| Phase angle | 10.26° | 26.11° | 89.33° | 346.26° | 325.90° | 294.52° | 104.74° | 346.62° |
| Vector length | 0.25 | 0.27 | 0.48 | 0.47 | 0.26 | 0.26 | 0.42 | 0.38 |
| Circular variance | 0.75 | 0.74 | 0.52 | 0.53 | 0.74 | 0.74 | 0.58 | 0.62 |
| Circular SD | 1.22 | 1.21 | 1.02 | 1.03 | 1.22 | 1.22 | 1.08 | 1.12 |
| Kurtosis | 0.08 | 0.02 | -0.02 | 0.21 | -0.06 | 0.12 | 0.10 | 0.07 |
| Skewness | -0.11 | 0.07 | 0.18 | 0.07 | -0.10 | 0.11 | 0.17 | -0.03 |

*Note.* Descriptive circular statistics, separated by age group, for the different conditions (pair and item), for the difference between them (pair – item), and for pair memory of the groups of high-performing individuals. Phase angle indicates the mean phase angle in degrees. The vector length indicates concentration around the mean phase angle. Circular variance and standard deviation (SD) are circular analogues to the corresponding linear measures. Kurtosis is a measure of peakedness and skewness indicates symmetry around the mean direction (i.e., mean phase angle).

**Table S2.** Descriptive and inferential statistics for covariate data.

|  | Younger adults |  | Older adults |  | Range | Wilcoxon-rank sum test |  |  |  |
| --- | --- | --- | --- | --- | --- | --- | --- | --- | --- |
| | M | SD | M | SD | | Rank sum | z | $\alpha$ | $p$ |
| DST | 66.24 | 10.79 | 49.91 | 10.14 |  | 4445 | 6.54 | .05 | <.001 |
| STW | 22.52 | 3.46 | 28.95 | 3.21 |  | 1989 | -7.58 | .05 | <.001 |
| MMST |  |  | 28.78 | 1.28 | 24–30 |  |  |  |  |

*Note.* Group means (M) and standard deviations (SD) are reported for each age group for the Digit Symbol substitution test (DST; 1) and the Spot-A-Word test (STW; 2), as well as inferential statistics for the group comparisons. The range for the Mini-Mental State Examination (MMST) is reported for older adults and.

**Table S3.**
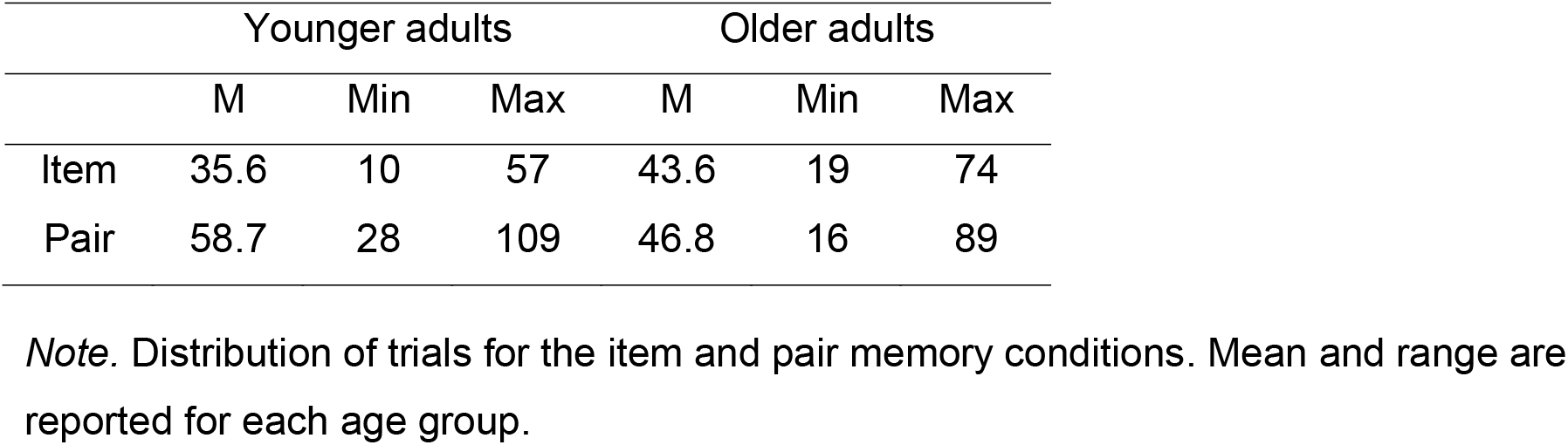
Distribution of trials per condition and age group.

## Notes

### Competing Interest Statement

The authors have declared no competing interest.

